# Turning movements in real life capture subtle longitudinal and preataxic changes in cerebellar ataxia

**DOI:** 10.1101/2021.03.22.436330

**Authors:** Annika Thierfelder, Jens Seemann, Natalie John, Martin Giese, Ludger Schöls, Dagmar Timmann, Matthis Synofzik, Winfried Ilg

**Author notes:** Corresponding authors: Winfried Ilg, Section, Computational Sensomotorics Hertie Institute for Clinical Brain Research, Otfried-Müller-Straße 25, 72076 Tübingen, Germany, or: Matthis Synofzik, Research Division Translational Genomics of Neurodegenerative Diseases, Center for Neurology & Hertie Institute for Clinical Brain Research, Hoppe-Seyler-Strasse 3, 72076 Tübingen, Germany. Authors have contributed equally (Annika Thierfelder, Jens Seemann). shared last authors (Matthis Synofzik, Winfried Ilg). Statistical analysis was conducted by Annika Thierfelder.

## Abstract

**OBJECTIVES:** Clinical and regulatory acceptance of upcoming molecular treatments in degenerative ataxias might greatly benefit from ecologically valid endpoints which capture change in ataxia severity in patients’ real life. This longitudinal study aimed to unravel quantitative motor biomarkers in degenerative ataxias in real life turning movements which are sensitive for changes both longitudinally and at the preataxic stage.

**METHODS:** Combined cross-sectional (n=30) and longitudinal (n=14, 1-year interval) observational study in degenerative cerebellar disease (including 8 pre-ataxic mutation carriers) compared to 23 healthy controls. Turning movements were assessed by three body-worn inertial sensors in three conditions: (1) instructed laboratory assessment, (2) supervised free walking, and (3) unsupervised real-life movements.

**RESULTS:** Measures which quantified dynamic balance during turning – lateral velocity change (LVC) and outward acceleration –, but not general turning measures such as speed, allowed differentiating ataxic against healthy subjects in real life with high effect size (δ=0.68), with LVC also differentiating preataxic against healthy subjects (δ=0.53). LVC was highly correlated with clinical ataxia severity (SARA score, effect size ρ=0.79) and subjective balance confidence (ABC score, ρ=0.66). Moreover, LVC in real life – but not general turning measures, gait measures, or the SARA score – allowed detecting significant longitudinal change in one-year follow-up with high effect size (r_prb_=0.66).

**CONCLUSIONS:** Measures of turning allow to capture specific changes of dynamic balance in degenerative ataxia in real life, with high sensitivity to longitudinal differences in ataxia severity and to the preataxic stage. They thus present promising ecologically valid motor biomarkers for capturing change in real life, even in the highly treatment-relevant early stages of degenerative cerebellar disease.

## Introduction

While manifold targeted molecular treatments for degenerative cerebellar diseases (DCDs) are now on the horizon^1, 2^, clinical and regulatory acceptance will depend on their proven effects on subject’s ataxia in real life. This highlights the need for quantitative ataxia biomarkers remotely monitored during subjects’ real life - ideally passively captured during usual everyday living. These quantitative-motor biomarkers should be sensitive to longitudinal change as well as to the early – possibly even preataxic – stages of ataxia disease, where molecular treatments are likely most effective^3, 4^.

Recent cross-sectional work focussing on the analysis of straight walking sequences has raised the possibility to capture quantitative motor changes in DCDs by remote sensor-based monitoring during daily life^5^. However, while measures of straight walking showed high sensitivity to cross-sectional ataxia severity^5^, other components of real-life walking behaviour like turning might place a higher coordinative motor demand and thus show a higher sensitivity to individual progression in particular for preataxic and early ataxia disease stages. This hypothesis receives support from our prior lab-based study showing that a coordinatively highly demanding task – tandem walking on a foam surface – revealed balance differences in the preataxic stage of DCD^6^. Turning movements in fact represent a highly relevant and coordinatively demanding type of everyday walking behaviour. As 35% to 45% of steps occur within turns^7^, they account for a substantial part of daily walking behaviour. Compared to straight walking, turning movements are also suggested to be more challenging in terms of dynamic balance^8-10^, as they involve a stronger demand of tuned anticipatory postural adjustments^11^ and specific trunk-limb coordination strategies^12^.

For the analysis of turning movements, existing work on Parkinson’s disease^13-17^, multiple sclerosis^18^, cerebellar ataxia^19, 20^, or elderly with recurrent falls^21, 22^ so far focused on the assessment of *general* turning parameters like turn angle, mean velocity, duration or the number of steps used to complete the turn. However, these measures do not aim to reflect specific dysfunctional control mechanisms like dynamic balance control. Such changes might be more sensitively captured by measures directly reflecting motor control mechanisms specifically impaired in cerebellar ataxias.

Based on these notions, we here hypothesized that dynamic balance measures of turning might be particularly sensitive to subtle ataxia changes not only under supervised task-based conditions, but also during unsupervised, task-free real life both (i) longitudinally and (ii) at preataxic and early stages of ataxia disease.

## Methods

### Participants

30 subjects at an ataxic or preataxic stage of degenerative cerebellar disease (DCD, age: 51 ±15 years) were recruited from the Ataxia Clinics of the University Hospitals Tübingen and Essen, Germany. They comprised of (i) 22 subjects at the ataxic stage of DCD (as defined by a Scale for the Assessment and Rating of Ataxia [SARA] score of ≥ 3 ^23^ (group ATX; mean SARA score of 9.4 points), and (ii) 8 subjects with repeat-expansions in SCA2, SCA 3 or SCA 6 at the preataxic stage of DCD (SARA score <3) (group PRE; mean SARA score of 1.37 points)^23^. For details of patient characteristics, see Table 1. DCD subjects were included based on following inclusion criteria: 1.) manifest or repeat expansion for DCD in the absence of any signs of secondary or other CNS disease; 2.) age between 18 and 75 years; 3.) ability to walk without walking aids. The exclusion criteria were: severe visual or hearing disturbances, cognitive impairment, predominant non-ataxia movement disorders (e.g. parkinsonism, spasticity), or orthopaedic constraints. 22 of the 30 DCD subjects (73.3%) carried a repeat expansion in SCA1, 2, 3 or 6. Severity of ataxia was rated using the SARA^23^. The SARA score includes the following 8 items: 3 items rating gait and posture, 1 item for speech disturbances, and 4 items for limb-kinetic functions. The 3 items rating gait and posture are grouped into the subscore SARA_posture&gait_ (SARA_p&g_)^6, 24^. In addition, we recruited 23 healthy controls (group HC: age: 48±15 years). HC subjects had no history of any neurological or psychiatric disease, no family history of neurodegenerative disease, and did not show any neurological signs upon clinical examination. Group sizes have been estimated based on earlier lab-based and real life ataxic gait studies^5, 6, 25-27^.

**Table 1.**
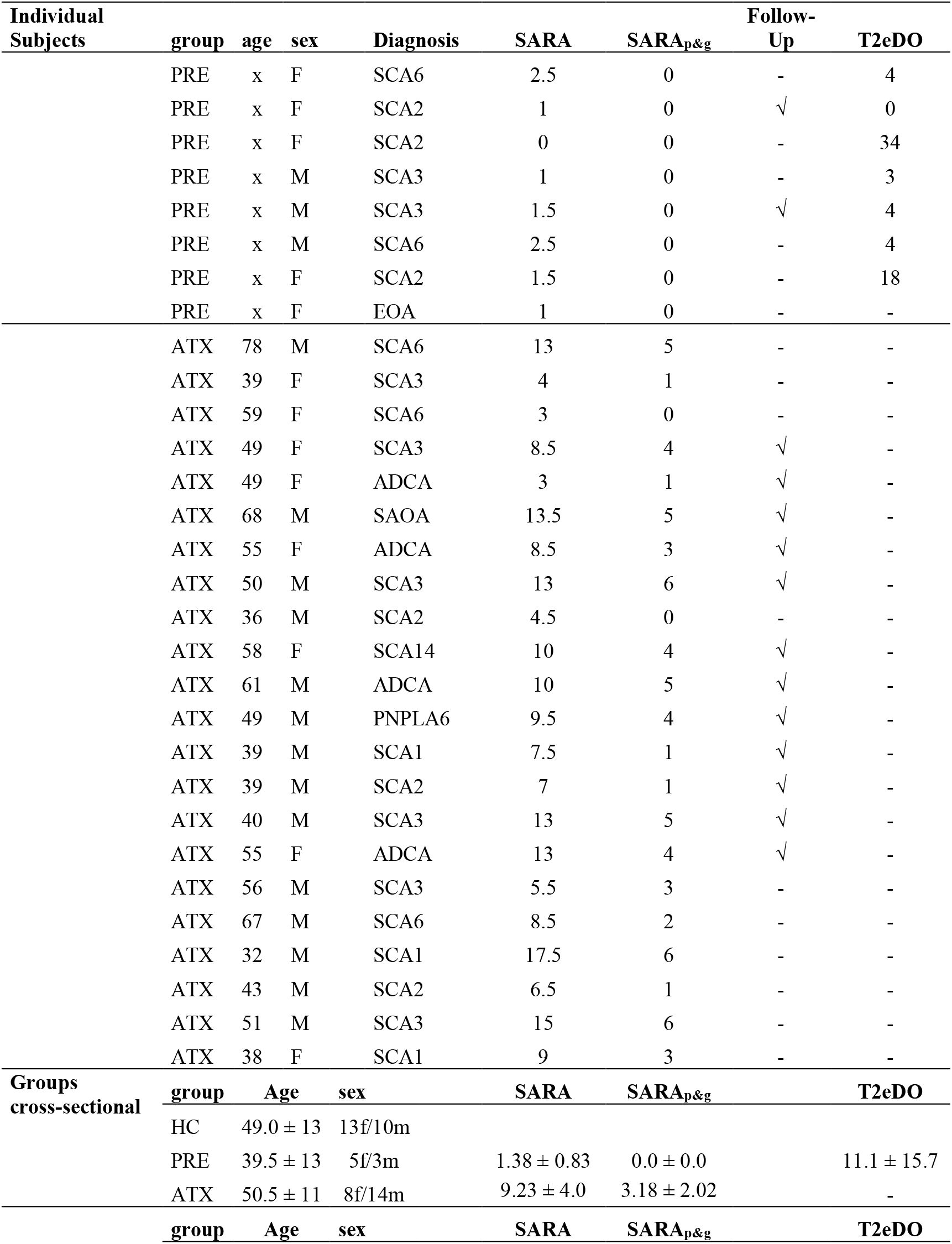
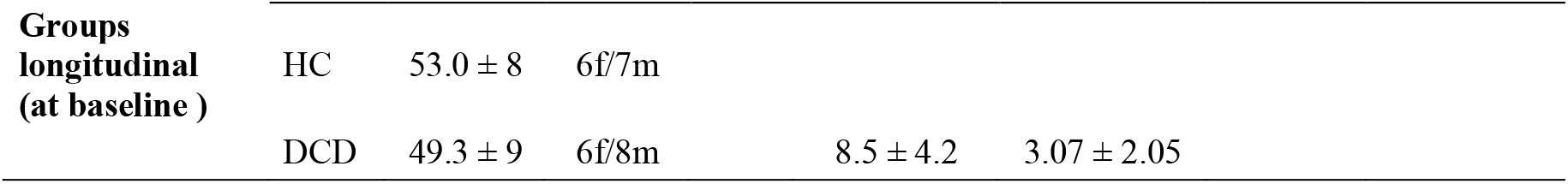
Subject characteristics. Clinical ataxia severity was determined by the SARA^23^. The SARA_posture&gait_ subscore is defined by the first three items of the SARA score which capture gait, standing and sitting ^24^. SAOA: sporadic adult onset ataxia; ADCA: autosomal dominant ataxia of still undefined genetic cause; EOA: early-onset ataxia of still undefined genetic cause. In this particular subject (PRE 8) with MRI evidence of cerebellar atrophy, yet still at the preataxic stage. SCA: autosomal-dominant spinocerebellar ataxia of defined genetic type. The following diagnoses denote the gene underlying the respective ataxia type: PNPLA6, ANO10 (=SCAR 10, autosomal-recessive spinocerebellar ataxia type 10). T2eDO: genetically estimated timespan to disease onset for preataxic SCA subjects (in years, calculated according to ^56^). At the bottom of the table, the average values for the different groups in the cross-sectional and the longitudinal analyses are provided: HC (healthy controls), PRE (preataxic subjects), ATX (ataxic subjects), DCD (degenerative cerebellar disease, comprising preataxic and ataxic subjects). For preataxic subjects we provide no individual age (x), since this would facilitate an individual identification of mutation carriers.

Longitudinal real life data from a one-year follow-up assessment (duration: 391±69 days) were recorded from 14 DCD subjects (12 ataxic, 2 preataxic) and 13 healthy controls. Reasons for longitudinal drop-out were unavailability for follow-up assessment (n=13), technical problems in follow-up assessment (n=1), and disability in walking without walking aids at follow-up (n=2). The study procedure was approved by the local ethics committee (598/2011BO1). All participants gave their informed consent prior to participation.

### Turning Conditions

Turning movements were recorded in three different conditions: two supervised conditions for validation purposes and one unsupervised condition in real life as the main target condition (For an overview, see Table 2): (i) **I**nstructed **T**ask-based **T**urning within a constrained turning task (ITT condition): Subjects were instructed to walk along a parkour at the T-junction of a lab corridor performing 90° and 180° turns as illustrated in Figure 1C, supervised by a study assessor. This task was conducted twice in each direction (complete walking time over all participants: 21.6± 8.1 s).

**Table 2.** Description of Turning conditions.

| **Assessment** | **Condition** | **Description** |
| --- | --- | --- |
| Task-based assessment | Lab-based turning based on an instructed standardized Turning Task (ITT) | Subjects were instructed to walk along the T-junction of a corridor performing 90° and 180° turns as illustrated in Figure 1C. |
| Supervised Free Assessment | Supervised Free Turning (SFT) | Subjects walked on public indoor and outdoor spaces on an institutional compound for 5 minutes at their preferred speed without pre-specified route, but supervised by a study assessor. Supervised free turnings (SFT) and supervised free walking bouts (SFW) were analysed separately. |
| Real Life Assessment | Real-Life Turning (RLT) | Subjects walked as part of their individual routine of daily living in their usual indoor and outdoor settings for 4-6 hours, without pre-specified routes or supervision. Real life turnings (RLT) and real life walking bouts (RLW) were analysed separately. |

**Figure 1.**
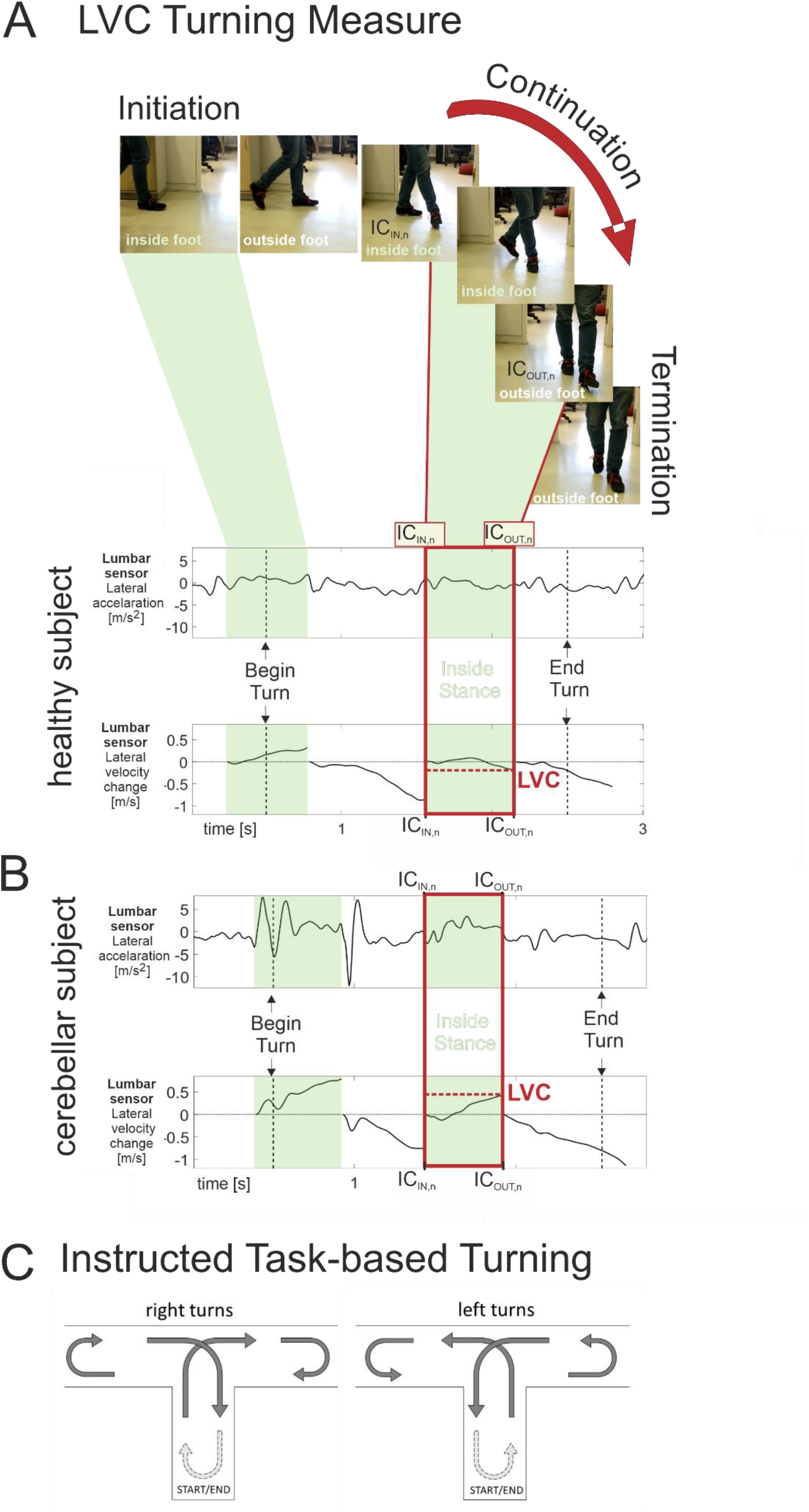
Illustration of the LVC (Lateral Velocity Change) turning measure and the instructed task-based turning walkway. (A) Computation of the LVC turning measure. Shown are snapshots of the steps of an exemplary 90° turn of a healthy subject and the corresponding trajectories of lateral acceleration and lateral velocity change of the lumbar sensor. The LVC measure is determined in inside stance phases (highlighted in green) during the continuation phase (highlighted by red boxes) of the turning movement. This phase lasts from the initial contact (IC) of the inside stance leg (IN) until the next initial contact of the outside (OUT) stance leg. The LVC measure is calculated by integrating the lateral acceleration in the described phase. (B)Corresponding diagrams of lateral acceleration and the LVC measure for a 90° turn of a cerebellar patient with moderate ataxia (SARA =10). (C) Schematic illustration of the T-junction walkway on a real corridor for the Instructed Task-based Turning (ITT) performing 90° and 180° turns. This procedure was generally conducted twice in each direction, resulting in eight 90° turns (four left and right turns, respectively), and ten 180° turns for each subject.

1. **S**upervised **F**ree **T**urning (SFT condition): Turning movements were extracted from unconstrained walking in public indoor and outdoor spaces on an institutional (hospital) compound where subjects were free to choose and change the floors and indoor and outdoor spaces where they wished to walk (complete walking time: 5 minutes). All spaces were open to public, but the subject’s walking performance was constantly supervised by a study assessor.
2. **R**eal **L**ife **T**urning (RLT condition): Turning movements were extracted from unconstrained walking during subjects’ everyday living without supervision by study personnel (total recording time per subject: 4-6 hours). Subjects were instructed to wear the sensors inside and outside their house, and to include at least a half-hour walk. Subjects documented their recorded walking movements in an activity protocol.

To capture the impact of disease on subjective confidence in activities of daily living, DCD subjects were asked to self-report their balance confidence using the Activity-specific Balance Confidence Scale (ABC)^28^ and two specific questions about the confidence in turning movements (see Supplementary Information).

### Methods for measuring turning movements

Three Opal inertial sensors (APDM, Inc., Portland, US) were attached to both feet, and to the posterior trunk at the level of L5 with elastic Velcro bands. Inertial sensor data was collected and wirelessly streamed to a laptop for automatic generation of gait and balance metrics by Mobility Lab software (APDM, Inc., Portland, US). For the turning conditions during unconstrained walking (SFT, RLT), data were logged on each OPAL sensor and downloaded after the session. Turns were identified by the Mobility Lab software, where the angular velocity about the vertical axis exceeded a threshold of 15°/ s. The start and end of each turn were set to the point where the angular velocity dropped below 5°/ s ^16^. For each detected turn, the following features were extracted using the Mobility Lab software: turn angle, duration, turn velocity, steps within turn, and raw accelerometer data of all three sensors^16, 29^. For determining the lateral acceleration, the sensor data was reoriented from the sensor body frame into a global reference frame using the orientation estimates provided by the Mobility Lab software^16^. This global reference frame was used to align the lateral axis of the lumbar acceleration orthogonal with respect to gravity. Within extracted turns, step events were determined by an algorithm based on continuous wavelet transform^30^.

Since we observed very few 180° U-turns in the unconstrained conditions SFT and RLT, we did not include 180° turns in the main analysis (for results of the 180° U-turns in ITT, see Supplement, Table 6). For the constrained ITT condition we analysed 90° turning movements: for the unconstrained conditions SFT and RLT we included turns in the range between 50° and 120°. In SFT and RLT, turns were only included into the analysis if two regular steps before and after the turn were detected. We analysed a mean of 78 ± 18 turning movements in the RLT condition over all participants.

### Measures of dynamic balance in turning movements

In addition to general turning parameters (i.e. turn angle, duration, mean velocity and number of steps) we focused on measures which allow quantifying impaired dynamic balance control while turning, in particular lateral sway. This was operationalized by the lateral acceleration of the lumbar sensor within turning movements. Previous work on wearable sensors has shown that such lateral acceleration is correlated to a dynamic stability criterion (margin of stability^31^) during walking and turning^32^. This dynamic stability criterion was defined in the medio-lateral dimension by regarding the lateral acceleration 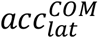 of the CoM (Centre of Mass) orthogonal to gravity and the direction of travel. Thus, the change in the lateral velocity velocity 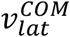 of the CoM during step *n* is given by

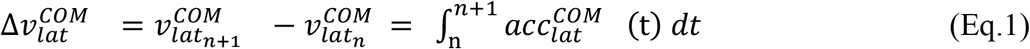

whereby 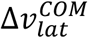 can be used to determine dynamic stability with respect to foot placement or to describe the amount of corrective foot placement needed to regain stability after a disturbance^31^.

Turning movements can be categorized in three phases: Initiation, Continuation, Termination (e.g. ^33^, see Figure 1A). Given that the largest whole-body angular momentums occur during the continuation phase^34^, our analysis focussed on the lateral acceleration during steps within the continuation phase in the middle of the turning movement, starting with the initial contact of the inside foot IC_IN_ until the subsequent initial contact of the outside foot (IC_OUT_) (see red boxes in Figure 1A+B).

The Lateral Velocity Change (LVC) of this period was computed by integrating the lateral acceleration (acc_lat_) of the lumbar sensor for step n and turning movement T, (Eq. 2).

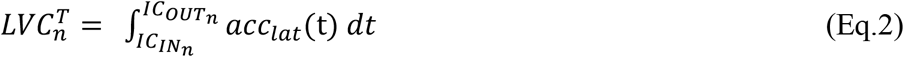

The LVC^T^ for one turning movement T was then determined by averaging all steps *n* LVC_n_^T^ within the turn. Note that for 90° turns there is often only one step which contributes to the LVC^T^ of that specific turning movement T (^34^, see Figure 1A+B). The resulting LVC measure for all turns of one subject in a condition was determined as the median of all LVC^T^ of corresponding turning movements. The LVC describes the relation between acceleration to the inside and outside of the turn curvature. To generalize across turns, we defined outward acceleration to be positive and inward acceleration to be negative. A positive LVC therefore denotes more velocity towards the outside of the turn curvature, while a negative LVC would point to more inward velocity.

As complementary measures, we also determined the amount of outward and inward acceleration separately, integrating absolute amounts of acceleration over the described periods in the continuation phase.

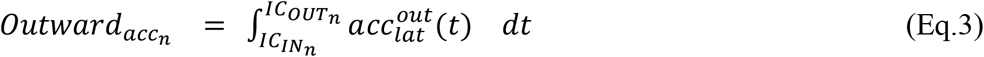

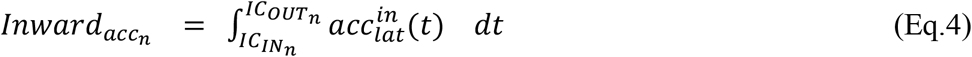

### Comparison of turning and gait measures in their sensitivity to preataxic and longitudinal changes

In addition, we examined whether *turning* measures show a higher sensitivity to early stage and longitudinal changes in DCD than *gait* measures. For this comparison, we selected those gait measures which had previously shown the highest effect sizes in sensitivity to ataxia severity in straight walking bouts in supervised as well as real life conditions, namely *lateral step deviation* and a *compound measure of spatial step variability* (combining *lateral step deviation* and *step length variability*)^5^. Gait parameters for free supervised (condition SFW) and real-life walking (condition RLW) were extracted from the same sessions as turning parameters, namely the trials of the conditions SFT and RLT. Gait parameters for lab-based gait assessment (condition LBW) were extracted from two 25 meters straight walking trials^5^. Gait parameters for all conditions are shown in the Supplement, Table 10.

### Statistics

Between-group differences (HC vs. ATX vs. PRE group) of movement measures were determined by the non-parametric Kruskal-Wallis-test. When the Kruskal-Wallis-test yielded a significant effect (p<0.05), post-hoc analysis was performed using a Mann-Whitney U-test. Effects sizes were determined by Cliff’s delta^35^.

Repeated measurements analyses were performed for the longitudinal analysis using the non-parametric Friedman test to determine within-group differences between assessments. When the Friedman test yielded a significant effect (p<0.05), post hoc analysis was performed using a Wilcoxon signed-rank test for pairwise comparisons between assessments. Effects sizes for the repeated measurements analyses were determined by matched pairs rank biserial correlation^36^.

We report three significance levels: (i) uncorrected *: p<0.05, (ii) Bonferroni-corrected for multiple comparisons **: p<0.05/n with n=3: number of analysed turning features of dynamic balance, (iii) ***p<0.001. Spearman’s ρ was used to examine the correlation between turning measures, gait measures, SARA scores and ABC scores. Effect sizes ρ were classified as ρ: 0.1 small effect, 0.3 medium effect, 0.5 large effect, 0.7 very large effect ^37, 38^. Statistical analysis was performed using MATLAB (Version R2018B). Based on the longitudinal changes of the LVC measure a sample size estimation was performed using G*power 3.1^39^ to determine the required cohort size for detecting a 50% reduction of natural progression by a hypothetical intervention.

## Results

### Group differences between HC, ATX and PRE for specific, but not general turning measures

No group differences were observed for general performance measures of turning including mean velocity, duration, turn angle and number of steps (Kruskal-Wallis test; Table 3). In contrast, the measures specifically reflecting dynamic balance in turning - namely LVC and outward acceleration - revealed significant group differences for all turning conditions. Post-hoc analysis revealed group difference between ATX and HC (p<0.006, Figure 2A and Table 3), with highest effect sizes (δ>0.68) in the RLT condition for both LVC and outward acceleration. Moreover, the LVC measure also revealed differences between PRE and HC, again with highest effect size for the RLT condition (p=0.029, δ=0.53, Figure 2A and Table 3).

**Table 3.** Cross-sectional analysis of turning measures. Between-group differences of healthy controls (HC), preataxic subjects (PRE) and ataxic subjects (ATX) for different turning measures in each of the three study conditions: Instructed Task-based Turning (ITT), Supervised free Turning (SFT) and Real Life Turning (RLT). Stars indicate significant differences between groups (*≡ p<0.05, **≡ p<0.01 Bonferroni-corrected, ***≡ p<0.001). KW denotes the result of the Kruskal-Wallis test. δ denote the effect sizes determined by Cliff’s delta. LVC: Lateral Velocity Change.

| Measure | Cond. | Descriptive Statistics |  |  | Statistical Test Results |  |  |  |  |  |  |
| --- | --- | --- | --- | --- | --- | --- | --- | --- | --- | --- | --- |
|  |  | HC | PRE | ATX | KW | HC vs PRE |  | HC vs ATX |  | PRE vs ATX |  |
| | | $\mu \pm \sigma$ | $\mu \pm \sigma$ | $\mu \pm \sigma$ | p | p | $\Delta$ | p | $\delta$ | p | $\delta$ |
| Mean Velocity | ITT | 53.59 ± 13.19 | 50.48 ± 8.77 | 50.89 ± 10.07 | 0.849 | 0.936 | 0.03 | 0.573 | 0.1 | 0.816 | 0.07 |
| | SFT | 45.81 ± 8.9 | 45.76 ± 5.58 | 47.8 ± 9.25 | 0.663 | 0.948 | 0.02 | 0.442 | 0.14 | 0.465 | 0.18 |
| | RLT | 46.68 ± 6.28 | 45.24 ± 3.76 | 47.98 ± 7.37 | 0.829 | 0.604 | 0.13 | 0.742 | 0.06 | 0.656 | 0.11 |
| Duration | ITT | 1.66 ± 0.3 | 1.71 ± 0.36 | 1.61 ± 0.33 | 0.821 | 0.851 | 0.06 | 0.523 | 0.11 | 1.00 | 0.01 |
| | SFT | 1.66 ± 0.34 | 1.59 ± 0.13 | 1.54 ± 0.31 | 0.729 | 0.528 | 0.15 | 0.521 | 0.12 | 0.724 | 0.09 |
| | RLT | 1.67 ± 0.26 | 1.67 ± 0.13 | 1.58 ± 0.3 | 0.566 | 0.718 | 0.09 | 0.286 | 0.19 | 0.76 | 0.08 |
| Angle | ITT | 84.63 ± 9.03 | 84.44 ± 5.63 | 80.09 ± 7.75 | 0.133 | 0.767 | 0.09 | 0.063 | 0.32 | 0.223 | 0.33 |
| | SFT | 74.03 ± 11.72 | 73.2 ± 7.92 | 69.35 ± 9.09 | 0.277 | 0.744 | 0.08 | 0.134 | 0.27 | 0.311 | 0.24 |
| | RLT | 73.61 ± 5.46 | 73.17 ± 3.47 | 70.01 ± 5.09 | 0.065 | 0.701 | 0.1 | 0.024* | 0.4 | 0.197 | 0.32 |
| #Steps | ITT | 3.48 ± 0.68 | 4.08 ± 1.02 | 3.35 ± 0.79 | 0.365 | 0.194 | 0.34 | 0.849 | 0.03 | 0.191 | 0.33 |
| | SFT | 3.39 ± 1.22 | 2.89 ± 0.74 | 2.83 ± 0.88 | 0.207 | 0.274 | 0.25 | 0.097 | 0.29 | 0.73 | 0.08 |
| | RLT | 3.04 ± 0.82 | 3.0 ± 0.53 | 2.7 ± 0.77 | 0.29 | 0.884 | 0.04 | 0.167 | 0.23 | 0.236 | 0.27 |
| LVC | ITT | -0.2 ± 0.19 | -0.19 ± 0.12 | -0.05 ± 0.25 | 0.104 | 0.809 | 0.07 | 0.052 | 0.33 | 0.169 | 0.38 |
| | SFT | 0.04 ± 0.2 | 0.06 ± 0.14 | 0.22 ± 0.31 | 0.002** | 0.648 | 0.11 | 0.001*** | 0.59 | 0.013* | 0.59 |
| | RLT | 0.07 ± 0.1 | 0.16 ± 0.07 | 0.26 ± 0.17 | <0.001*** | 0.029* | 0.53 | <0.001*** | 0.68 | 0.214 | 0.31 |
| Outward<sub>Acc</sub> | ITT | 31.0 ± 13.5 | 27.0 ± 7.4 | 47.2 ± 24.0 | 0.006** | 0.319 | 0.28 | 0.006** | 0.47 | 0.021* | 0.63 |
| | SFT | 46.5 ± 14.4 | 44.97 ± 12.17 | 61.8 ± 13.0 | 0.003** | 0.948 | 0.02 | 0.002** | 0.59 | 0.011* | 0.62 |
| | RLT | 44.5 ± 8.7 | 49.7 ± 8.2 | 63.9 ± 17.4 | <0.001*** | 0.13 | 0.37 | <0.001*** | 0.7 | 0.037* | 0.51 |
| Inward<sub>Acc</sub> | ITT | 57.84 ± 19.34 | 52.51 ± 18.07 | 54.5 ± 21.56 | 0.541 | 0.27 | 0.304 | 0.489 | 0.12 | 0.697 | 0.11 |
| | SFT | 42.87 ± 16.12 | 35.04 ± 8.81 | 28.83 ± 9.57 | 0.008** | 0.157 | 0.33 | 0.003** | 0.55 | 0.129 | 0.37 |
| | RLT | 34.3 ± 10.41 | 28 ± 3.58 | 29.93 ± 8.56 | 0.18 | 0.075 | 0.435 | 0.192 | 0.229 | 0.725 | 0.09 |

**Figure 2.**
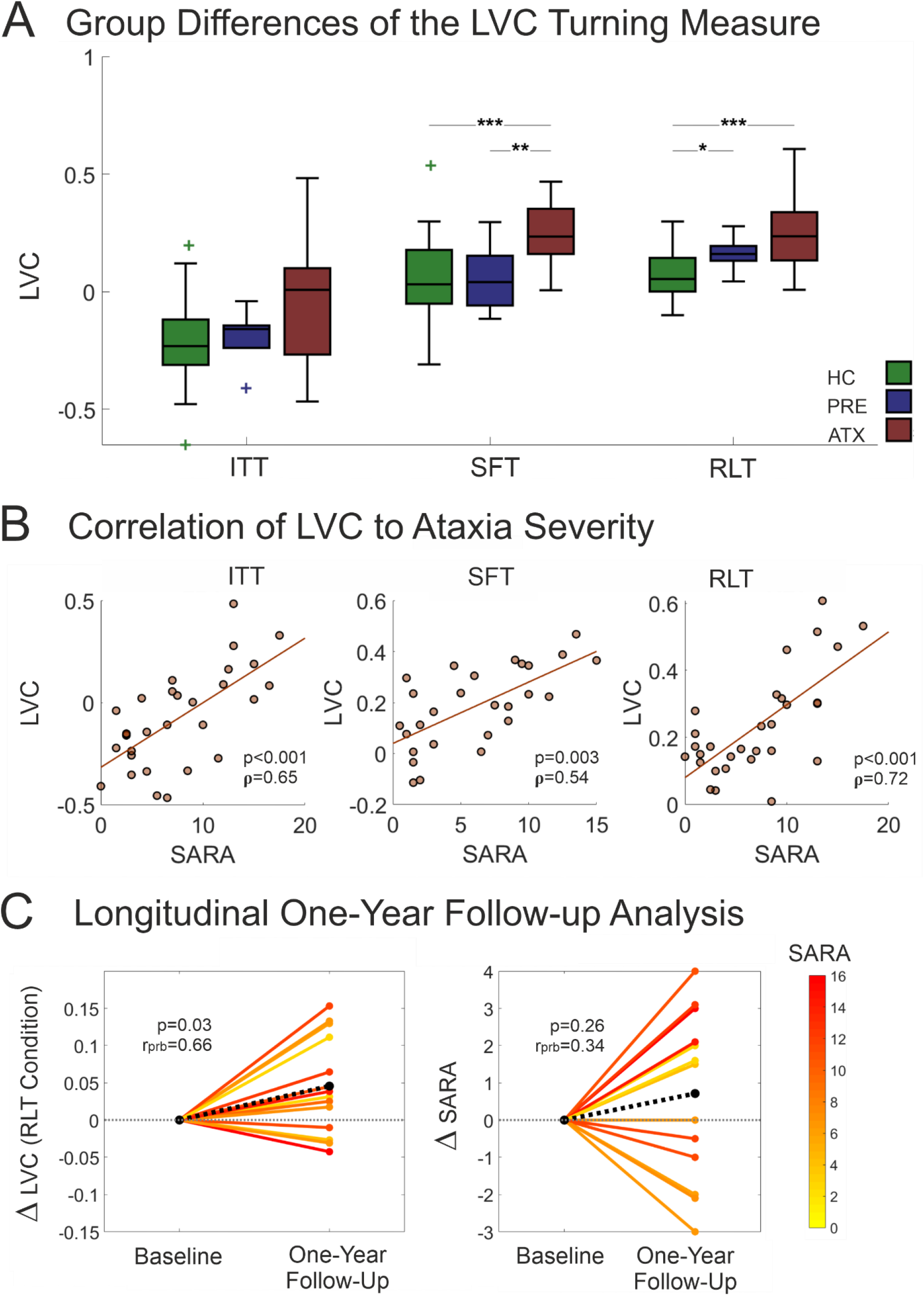
Group differences, relation to clinical ataxia severity and change over time for the LVC turning measure. (A) Between-group differences of the LVC measure for healthy controls (HC, green), preataxic subjects (PRE, blue) and ataxic subjects (ATX, red) in the different turning conditions: Instructed Task-based Turning (ITT), Supervised Free Turning (SFT) and Real Life Turning (RLT). (B) Relationship between SARA score and the LVC measure for the different turning conditions. Shown are all subjects with degenerative cerebellar disease (DCD) including both preataxic and ataxic subjects. (C) Within-subject changes between baseline and one-year follow-up for cerebellar subjects (DCD): (Left panel) Within-subject changes of the measure LVC in the real life turning condition RLT. (Right panel) Within-subject changes of the SARA score. In both panels SARA scores of cerebellar subjects are colour coded. Black dotted line = mean change across all subjects. Stars indicate significant differences between groups (*≡ p<0.05, **≡ p<0.01 Bonferroni-corrected, ***≡ p<0.001).

### Sensitivity of turning measures to ataxia severity: cross-sectional analysis

To analyse sensitivity of the turning measures to ataxia severity, we correlated these measures with the clinical ataxia score SARA and the SARA_posture&gait_ (see Methods) for the ATX group (as the SARA reflects clinical ataxia severity only for ataxic, not preataxic subjects^6, 40^). Out of the general turning measures, the mean velocity and number of steps during turning showed correlations only for the SFT condition (Table 4). In contrast, the measures specifically reflecting dynamic balance while turning - LVC and Outward_Acc_ - revealed highly significant correlations to the SARA score across all turning conditions, with again highest effect size (ρ=0.79) for real life turning movements (Table 4, see Figure 2B for correlations including the preataxic subjects).

**Table 4.** Correlations of turning measures with clinical and patient-oriented scores. Correlations between turning measures and clinical ataxia severity (SARA^23^ total score and SARAp&g posture&gait subscore^24^) as well as subjects’ self-reported subjective confidence ratings of balance in everyday activities (ABC-score)^28^. In addition to the validated questions of the ABC score, we asked the patients two specific questions about their balance confidence in everyday life turning movements of 90° and 180° (see supplementary Information). Correlations were determined for the cohort of ataxic subjects (ATX) in the three study conditions: Instructed Task-based Turning (ITT), Supervised free Turning (SFT) and Real Life Turning (RLT). LVC: Lateral Velocity Change. (*≡ p<0.05, **≡ p<0.01 Bonferroni-corrected, ***≡ p<0.001). Effect sizes of correlations are given using Spearman’s ρ.

| Turning measure | Condition | SARA |  | SARA<sub>p&g</sub> |  | ABC score |  | Question 90° Turns |  | Question 180° Turns |  |
| --- | --- | --- | --- | --- | --- | --- | --- | --- | --- | --- | --- |
| | | p | $\rho$ | p | $\rho$ | p | $\rho$ | p | $\rho$ | p | $\rho$ |
| LVC | ITT | <0.001*** | 0.691 | <0.001*** | 0.727 | <0.001*** | -0.51 | 0.002** | -0.46 | 0.016* | -0.36 |
|  | SFT | 0.005** | 0.628 | 0.027* | 0.521 | 0.015* | -0.6 | 0.002** | -0.72 | 0.083 | -0.45 |
|  | RLT | <0.001*** | 0.789 | <0.001*** | 0.734 | <0.001*** | 0.66 | 0.026* | 0.45 | 0.058 | 0.39 |
| Outward<sub>Acc</sub> | ITT | 0.003** | 0.585 | 0.002** | 0.611 | 0.018* | -0.36 | 0.024* | -0.34 | 0.093 | -0.26 |
|  | SFT | 0.017* | 0.554 | 0.028* | 0.518 | 0.038* | -0.52 | 0.01* | -0.62 | 0.154 | -0.37 |
|  | RLT | <0.001*** | 0.816 | <0.001*** | 0.747 | 0.009** | 0.52 | 0.094 | 0.35 | 0.206 | 0.27 |
| Inward<sub>Acc</sub> | ITT | 0.083 | 0.361 | 0.081 | 0.363 | 0.031* | 0.33 | 0.09 | 0.26 | 0.159 | 0.22 |
|  | SFT | 0.199 | 0.317 | 0.164 | 0.342 | 0.039* | 0.52 | 0.001*** | 0.78 | 0.06 | 0.48 |
|  | RLT | 0.127 | 0.336 | 0.212 | 0.277 | 0.109 | 0.34 | 0.204 | 0.27 | 0.11 | 0.33 |
| Mean Velocity | ITT | 0.238 | 0.25 | 0.252 | 0.243 | 0.89 | -0.02 | 0.888 | 0.022 | 0.893 | 0.021 |
|  | SFT | 0.001** | 0.699 | 0.001** | 0.732 | 0.352 | -0.25 | 0.09 | -0.44 | 0.589 | -0.15 |
|  | RLT | 0.622 | 0.111 | 0.332 | 0.217 | 0.733 | 0.074 | 0.517 | 0.14 | 0.325 | 0.21 |
| Duration | ITT | 0.567 | 0.123 | 0.343 | 0.202 | 0.808 | 0.039 | 0.834 | -0.03 | 0.916 | 0.017 |
|  | SFT | 0.317 | 0.25 | 0.096 | 0.405 | 0.003** | 0.68 | 0.024* | 0.56 | 0.002** | 0.72 |
|  | RLT | 0.49 | 0.155 | 0.284 | 0.239 | 0.38 | 0.19 | 0.754 | 0.068 | 0.211 | 0.26 |
| Angle | ITT | 0.088 | 0.356 | 0.17 | 0.289 | 0.379 | 0.14 | 0.269 | 0.17 | 0.335 | 0.15 |
|  | SFT | 0.089 | 0.412 | 0.224 | 0.301 | 0.05 | 0.5 | 0.181 | 0.35 | 0.006** | 0.66 |
|  | RLT | 0.131 | 0.332 | 0.422 | 0.18 | 0.627 | 0.1 | 0.348 | 0.2 | 0.537 | 0.13 |
| #Steps | ITT | 0.665 | 0.093 | 0.768 | 0.064 | 0.176 | 0.21 | 0.273 | 0.17 | 0.354 | 0.14 |
|  | SFT | 0.014* | 0.57 | 0.005** | 0.635 | 0.068 | 0.47 | 0.002** | 0.71 | 0.133 | 0.39 |
|  | RLT | 0.713 | 0.083 | 0.878 | 0.035 | 0.771 | 0.063 | 0.585 | 0.12 | 0.381 | 0.19 |

In addition to the correlation with clinical ataxia severity (SARA), the LVC measure also revealed high correlations with the subjective balance confidence in activities of daily living, assessed by the ABC score (p<0.001, effect size ρ=0.66, Table 4).

### Sensitivity of turning measures, but not gait measures, to longitudinal change in real life

We next analysed whether the turning measures allow to detect longitudinal changes in real life in DCD subjects at one-year follow-up assessment. While the clinical scores SARA (baseline mean: 8.5, follow-up mean: 9.2, p=0.26, effect size r_prb_=0.34) and SARA_p&g_ as well as general turning measures failed to detect significant longitudinal changes (Table 5 and Figure 2C), paired statistics revealed significant differences between baseline and one-year follow-up for the specific turning measure LVC (p=0.03, r_prb_=0.66; Table 5 and Figure 2C). The longitudinal increase of the LVC measure indicates a more pronounced acceleration in the outward direction of around 21% of the difference between HC and DCD at baseline. Sample size estimation shows a required cohort size of n=66 for detecting a 50% reduction of natural progression by a hypothetical intervention (80% power and one-sided 5% type I error).

**Table 5.** Longitudinal analysis of turning and gait measures. Longitudinal within-subject comparison of clinical ataxia ratings (SARA^23^ total score and SARAp&g posture&gait subscore^24^) as well as turning and gait measures in real life for baseline and 1-year-follow-up (p-values determined by Wilcoxon signed-rank test; effect sizes rprb determined by matched-pairs rank-biserial correlation ^36^). Shown are analyses for healthy controls (HC) and the group of degenerative cerebellar disease (DCD), consisting of preataxic and ataxic subjects. LVC: Lateral Velocity Change. Stars indicate significant differences between groups (*≡ p<0.05, **≡ p<0.016 Bonferroni-corrected, ***≡ p<0.001)

| Assessment Type | Measure | Group | Descriptive Statistics |  | Statistical Testing |  |
| --- | --- | --- | --- | --- | --- | --- |
| | | | Baseline | Follow Up | p | $r_{prb}$ |
| Clinical Measures | SARA | DCD | 8.5 ± 4.2 | 9.2 ± 4.3 | 0.26 | 0.34 |
| | SARA<sub>p&g</sub> | DCD | 3.1 ± 2.1 | 3.3 ± 1.9 | 0.438 | 0.35 |
| Turning Measures - Real Life | Angle | DCD | 70 ± 6.1 | 69 ± 3.2 | 0.855 | 0.07 |
| | | HC | 72 ± 5.7 | 73 ± 8 | 0.946 | 0.03 |
| | Mean Velocity | DCD | 49 ± 8.5 | 49 ± 8.4 | 0.67 | 0.14 |
| | | HC | 45 ± 6.6 | 44 ± 8 | 0.34 | 0.32 |
| | #Steps | DCD | 2.6 ± 0.74 | 2.7 ± 0.72 | 1.00 | 0.1 |
| | | HC | 3.1 ± 0.95 | 3.2 ± 0.69 | 0.883 | 0.04 |
| | Duration | DCD | 1.5 ± 0.33 | 1.5 ± 0.31 | 0.625 | 0.16 |
| | | HC | 1.7 ± 0.29 | 1.7 ± 0.32 | 0.508 | 0.22 |
| | LVC | DCD | 0.27 ± 0.17 | 0.31 ± 0.17 | 0.03 * | 0.66 |
| | | HC | 0.059 ± 0.1 | 0.047 ± 0.14 | 0.588 | 0.19 |
| | Outward<sub>Acc</sub> | DCD | 66 ± 17 | 69 ± 16 | 0.463 | 0.24 |
| | | HC | 45 ± 7.3 | 45 ± 10 | 0.542 | 0.21 |
| | Inward<sub>Acc</sub> | DCD | 33 ± 8.4 | 28 ± 8.7 | 0.049 * | 0.6 |
| | | HC | 37 ± 12 | 38 ± 16 | 0.588 | 0.19 |
| Gait measures - Real Life | Lateral Step Deviation | DCD | 4.4 ± 0.76 | 4.4 ± 0.74 | 0.715 | 0.12 |
| | | HC | 3.8 ± 0.25 | 3.8 ± 0.4 | 0.578 | 0.39 |
| | Compound Spatial Variability | DCD | 0.33 ± 0.13 | 0.32 ± 0.19 | 0.715 | 0.12 |
| | | HC | 0.19 ± 0.1 | 0.18 ± 0.07 | 0.898 | 0.06 |
| | Gait Speed | DCD | 1.1 ± 0.21 | 1.1 ± 0.28 | 0.194 | 0.41 |
| | | HC | 1.2 ± 0.19 | 1.2 ± 0.13 | 0.7 | 0.15 |

In contrast, there were no longitudinal changes in the LVC for the HC group (p>0.5), neither showed the gait measures any significant longitudinal change for the DCD or HC groups (Table 5).

## Discussion

Here we aimed to identify quantitative motor biomarkers for DCDs sensitive to subtle ataxia changes not only under supervised conditions, but also during real life by remote recording via wearable sensors, and with high sensitivity to both longitudinal change and the preataxic stages of disease. As turning movements are particularly challenging for dynamic balance control^9^, we hypothesized that turning measures specifically capturing dynamic balance control might be most sensitive for such ataxia-related movement changes in real life. Lateral velocity change (LVC), a novel measure specifically capturing dynamic balance, allowed differentiating not only ataxic subjects, but also preataxic subjects with high effect size in real life turning movements. Moreover, in contrast to general turning measures, gait measures, and the clinical SARA score, this specific measure allowed detecting significant longitudinal changes in one-year follow-up recordings, with likewise high effect size.

### Features of dynamic balance as a sensitive feature of ataxic turning movements

Compared to other turning features, the changes in dynamic balance while turning reflected by the LVC measure delivered the highest effect sizes in ataxic versus healthy subjects in all three conditions (Table 3). The specificity of LVC in capturing *ataxia-related* changes in dynamic balance control while turning is supported by the findings that (i), in contrast to LVC, no group differences were observed for ataxic versus control subjects in general turning measures like turn angle, mean velocity or number of steps (Table 3), and (ii) no correlation were found between LVC and general turning measures (Table 4). This indicates that the observed differences in LVC are not just secondary to general changes in turning or to different turning strategies^12^, which would also result in a change of other turning measures, e.g. higher velocity, differences in turn angles or a decreased number of steps. So far, ataxic turning movements have been described with such general movement parameters, resulting in characterizations such as enlarged base of support, shortened step length, and increased number of steps^20^. Most likely, these changes mainly reflect a set of *compensatory* strategies which are aiming at reducing the instability arising with turning^20^. Such compensation-induced movement changes for avoiding instabilities and falls are probably more pronounced in stages of more advanced ataxia severity. They are thus likely less relevant in the early stages of DCD, as they do not directly capture changes in cerebellar-related mechanisms like dynamic balance control. Our novel measures may thus instead provide a more direct characterization (and quantification) of ataxia-related changes in dynamic balance while turning,

Comparing the measures of dynamic balance across the different turning conditions, we observed high correlations between the constrained lab- and task-based (ITT) and the unconstrained task-free (SFT, RLT) turning conditions, in particular for LVC (p<0.001, ρ>0.64, Table 8). This is notable because of two aspects: First, turning behaviour during unconstrained walking is more variable compared to standardized task-based assessments; Second, we considered turning movements in the range between 50° and 120° for the free walking conditions, as these naturally occur with highest frequency during unconstrained walking, while only 90° turns were analysed in the standardized task-based assessment. The correlations between conditions suggest that our turning measure validly captures characteristics of turning movements in real-life behaviour, as it is validated in strictly standardized and supervised turns. Moreover, they indicate that also standardized assessments can be exploited to deliver first surrogate snapshots of patients’ unconstrained turning performance. However, this might come at a cost of partly less ecological validity and in particular smaller effect sizes: Effect sizes in group differences and correlations were highest in the unconstrained real life condition (Table 3 and 4), most probably due to the larger amount of turns in this condition. This finding highlights the power of remote movement capture by means of task-free, passive monitoring of subject’s ataxia severity during patient’s usual daily living. At the same time, it might point to a possible limitation of task-based remote movement capture which samples only short snapshots of standardized surrogate tasks for ataxia behaviour, as e.g. in instrumented SARA assessments ^41^.

**Table 6.**
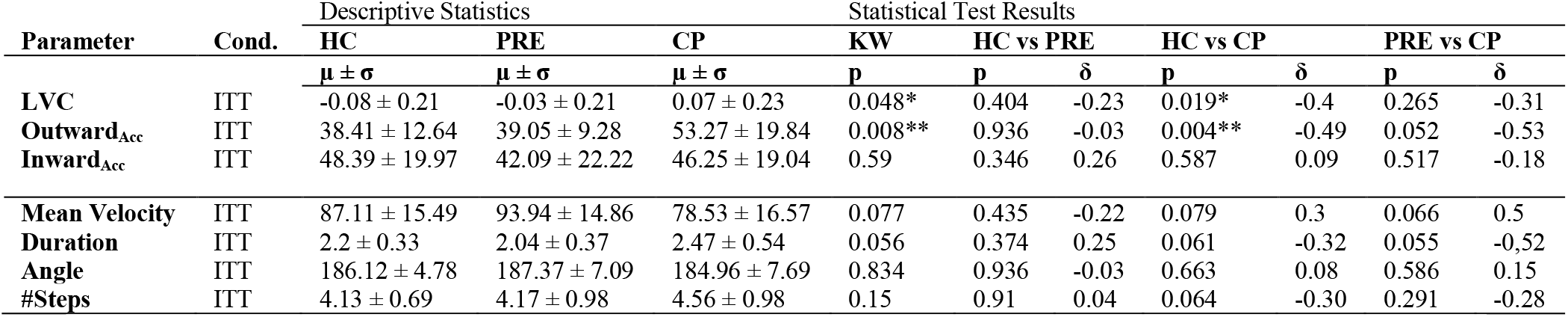
Analysis of 180° turns. Between-group differences of healthy controls (HC), preataxic subjects (PRE) and ataxic subjects (ATX) in turning measures for 180° U-Turns, shown for the lab-based turning condition ITT. Stars indicate significant differences between groups (*≡ p<0.05, **≡ p<0.01 Bonferroni-corrected, ***≡ p<0.001). KW denotes the result of the Kruskal-Wallis test. δ denotes the effect sizes determined by Cliff’s delta.

**Table 7.**
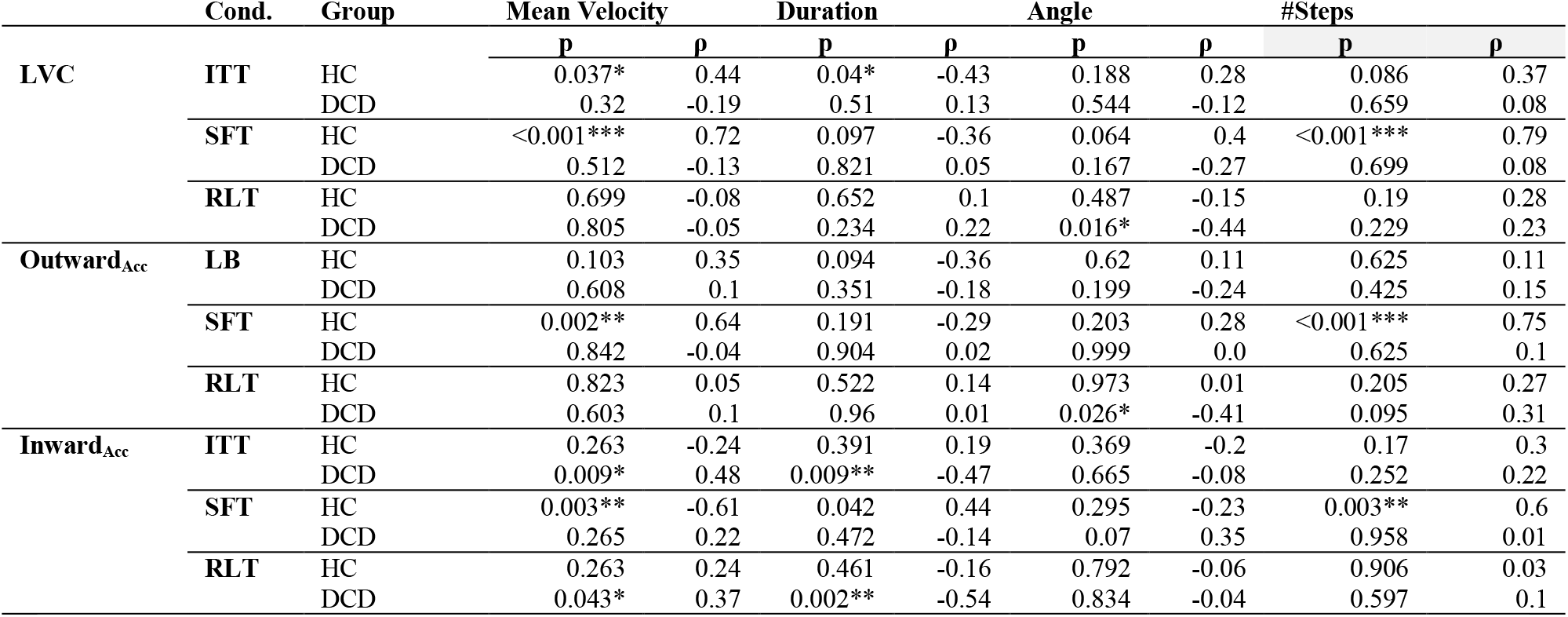
Correlations between turning measures related to dynamic balance and general turning measures. Shown are the correlations for healthy controls (HC) and for subjects with degenerative cerebellar disease (DCD, consisting of both preataxic and ataxic subjects) in the different turning condition: Instructed Task-based Turning (ITT), Supervised Free Turning (SFT) and Real Life Turning (RLT). (*≡ p<0.05, **≡ p<0.01 Bonferroni-corrected, ***≡ p<0.001). Effect sizes of correlations are given using Spearman’s ρ. LVC: Lateral Velocity Change.

**Table 8.**
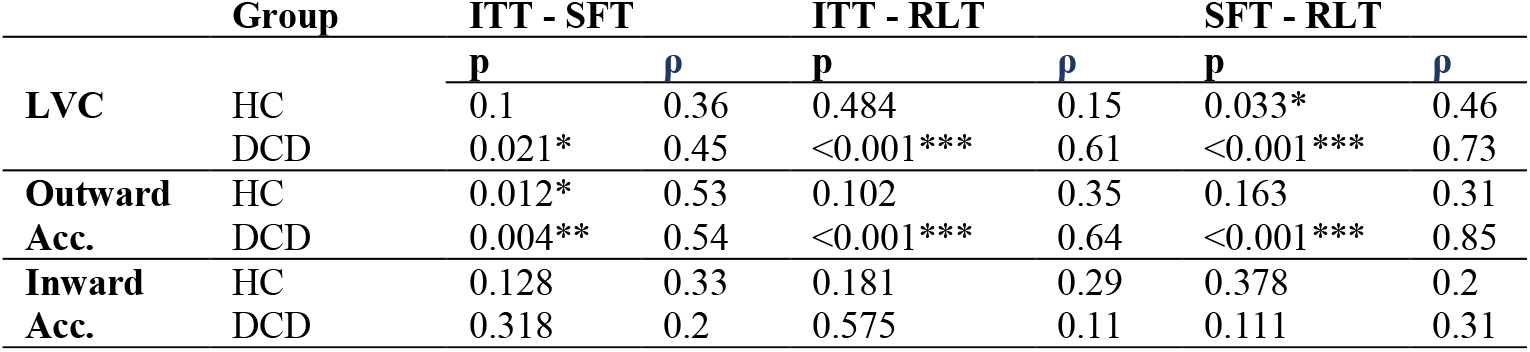
Correlation of turning measures of dynamic balance between turning conditions. Instructed Task-based Turning (ITT), Supervised Free Turning (SFT) and Real Life Turning (RLT). Given are correlation for healthy subjects (HC) as well as for subjects with degenerative cerebellar disease (DCD, consisting of both preataxic and ataxic subjects). (*≡ p<0.05, **≡ p<0.01 Bonferroni-corrected, ***≡ p<0.001). Effect sizes of correlations are given using Spearman’s ρ. LVC: Lateral Velocity Change.

### Measures of ataxic turning during real life are sensitive to ataxia severity and correlate with self-reported balance confidence in cross-sectional analyses

The measures capturing dynamic balance while turning - *LVC* and *Outward*_*acc*_ - were highly correlated to clinical ataxia severity in all conditions, with highest effect sizes in real life (see Table 4, Figure 2B). In contrast, general turning measures like mean velocity, turning angle and number of steps failed to reach significance correlations to ataxia severity in real life.

In addition, our digital-motor measures reflecting dynamic balance also correlated with the subjects’ self-reported confidence in real life balance activities, as quantified by the ABC-score (Table 4). Taken together, this close correlation with both a clinician-reported outcome (SARA) and a patient-reported outcome (ABC) indicates the validity of our measure as a real-life digital-motor biomarker for clinical trials. Specifically, it emphasizes the functional validity of our turning measures, affirming that they reflect overall ataxia severity and real life balance confidence. It is also consistent with an earlier study of multiple sclerosis (MS) which identified turning as an important marker of balance confidence and walking limitations^42^.

### Measures of ataxic turning during real life are sensitive to the preataxic stage

In addition to the differentiation of ataxic patients from healthy controls, our results are the first to show for the first time a group difference also between preataxic subjects and healthy controls in real life walking behaviour. The preataxic stage of SCAs attracts increasing research interest as it could provide a promising window for early therapeutic intervention - both pharmaceutical and rehabilitative - before substantial irreversible neurodegeneration has occurred^1, 3, 4^.

The observation that we found preataxic changes in turning movements (Table 3), but not in straight walking (Supplement Table 10), supports the hypothesis that turning is more challenging in terms of dynamic balance. This is also consistent with our earlier study on preataxic subjects which identified changes in a coordinatively more demanding walking type - tandem walk on a mattress - but not in straight walking^6^.

However, there is some inconsistency in the literature, with other studies having reported pre-ataxic changes in straight walking during clinical gait assessments^43, 44^. This discrepancy might most likely be explained with early clinical gait signs already present in these study cohorts ^43,44^. None of the preataxic subjects in our cohort showed any clinical gait or balance sign, as indicated by a SARA_posture&gait_ subscore = 0 for all preataxic subjects (Table 1).

### Measures of ataxic turning are sensitive to longitudinal change in real life

To serve as progression and treatment outcome measures, measures of real-life walking behaviour should ideally also be able to capture longitudinal changes that correspond to clinically important differences and relevant changes in patient-centred outcome measures ^1, 45^. Longitudinal progression studies in DCDs are still rare, and largely limited to clinical and imaging outcome measures^46-51^. In a large European multi-centre longitudinal study, mean annual SARA progression rates range from 2.11 points per year for SCA 1 to 0.8 points per year for SCA6 ^48^ and have been suggested to be even slower for non-repeat SCAs^52, 53^. Only two studies examined the longitudinal course of quantitative gait measures ^54, 55^: A study of a mixed cohort of 12 patients with DCD observed a significant increase in the SARA score at both the 2- and 4-year follow-up, while all the gait variables (with exception of an increased trunk rotation magnitude) showed a significant change only at the 4-year follow-up, indicating a rather limited sensitivity to change^54^. The other study on 11 degenerative ataxia subjects found a change in gait measures after one year only for normalized stance time in fast gait^55^.

In line with these previously reported progression rates of the clinical SARA score, we here observed a mean increase of the SARA score of 0.7 at one-year follow-up, which did not reach significance compared to baseline (p=0.26) (Table 5). Similarly, also the quantitative gait changes did not reach significance at one year-follow-up. In contrast, we observed significant changes in our measures capturing dynamic balance control in real life turning which again supports the notion that turning movements and the specific measures capturing its balance control component are sensitive to subtle changes. Given that we observed significant changes in a rather small study cohort (follow-up data available of n=14 subjects) with high effect size indicates that our measures might be very sensitive not only for longitudinal change, but likely also for treatment-related change in upcoming intervention trials. Sample size estimation revealed a required cohort size of n=66 for detecting a 50% reduction of natural progression by a hypothetical intervention. This seems to be remarkable as our study cohort also included rather slow progressive DCD types, e.g. SCA6^46^ and non-repeat SCAs^52^. In comparison, for clinical measures like SARA, earlier studies reported required cohort sizes of n>100 for detecting a 50% reduction of natural progression ^49^.

### Measures of ataxic turning seem particular sensitive at the early stages of degenerative ataxia disease

The measure LVC seems to be particularly sensitive for changes in balance control in early stages of DCD. This is indicated by significant difference between preataxic subjects versus healthy controls (Figure 2A, Table 3) as well as by a more detailed inspection of the longitudinal analysis. These data revealed that the longitudinal change observed for LVC might be driven by mild to moderate affected patients, whereas patients with more advanced ataxia (SARA ≥10 at baseline) showed no further change in the follow-up (Table 5 and Figure 2C). For patients in this more advanced disease stage, measures of straight walking might be more sensitive to change (but likely only until walking aids become necessary, when common gait measures then completely lose their sensitivity). Taken together, these findings indicate a disease-stage dependency of the effect sizes of digital-motor biomarkers in DCDs. This is highly important to be taken into account when now selecting the most appropriate outcome measures relative to the respective SARA-stratified inclusion criteria in upcoming targeted treatment trial in DCDs.

### Conclusions, limitations, and outlook

This study unravels measures reflecting dynamic balance control that allow to quantify real-life ataxic turning movements with high sensitivity to subtle changes in both (i) preataxic subjects as well as (ii) longitudinal progression in one-year follow-up. The findings are limited by the fact that our study cohort is not sufficiently powered for stratification according to specific ataxia genotypes and for detecting longitudinal change within the preataxic group only. Larger multi-centric future studies focussing on remote movement capture of real life behaviour using wearable sensors are required to demonstrate these possible promises of our measures.

However, our study might already have prepared first steps towards developing regulatory approval of digital-motor biomarkers as endpoints for future treatment trials in DCDs, by demonstrating (i) their power as ecologically valid biomarkers by capturing ataxia-specific motor behaviour in real life, (ii) their correlation with both clinical ataxia severity and self-reported balance confidence outcomes in real life, and (iii) their sensitivity to subtle changes longitudinally and at the earliest stages of the disease. It will be these early disease stages of DCD which are crucially important for upcoming molecular treatment trials that aim to prevent disease progression^1, 3^. As our turning measures correlate with both clinical and patient-reported outcomes at the *ataxic* stage of DCD, a change to the natural history of these measures already at the *preataxic* stage (e.g. by disease-modifying treatments) might possibly be accepted as a surrogate endpoint also in treatment trials targeting *the preataxic stage*.

## Acknowledgements

This work was supported via the European Union’s Horizon 2020 research and innovation program under the frame of EJP-RD network PROSPAX (No 441409627; M.S., and D.T. as an associated partner), and in part, by the German Hereditary Ataxia Society (DHAG), the “Stiftung Hoffnung” (to M.S.). Additional support has been received from BMG (project SStepKiZ to M.G) and an European Union’s ERC SNERGY Grant (RELEVANCE to M.G.). The authors thank the International Max Planck Research School for Intelligent Systems (IMPRS-IS) for supporting A.T and J.S..

## Supplementary Information

### Relationships between turning and gait measures

In order to relate disturbances in turning movements to gait abnormalities, we correlated the novel turning measures quantifying dynamic balance to those gait measures which had previously shown high sensitivity to ataxia severity in lab-based and real life conditions ^5^. The strongest correlations were observed between LVC and the compound gait measure of spatial variability for all three conditions, with the highest effect size in the real life condition (effect size ρ=0.61, see Table 9).

**Table 9.**
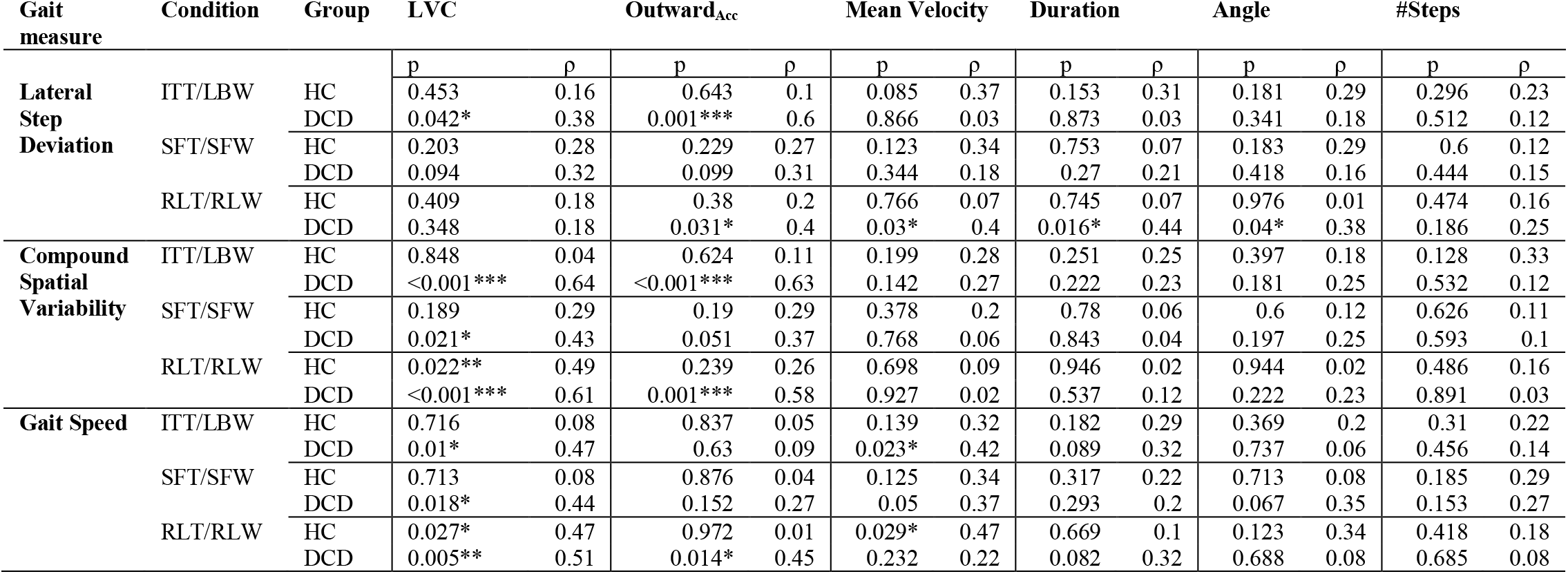
Relation between turning and gait measures. Correlations between turning and gait measures from the corresponding study conditions (ITT/LBW: Instructed lab-based turning/walking; SFT/SFW: S supervised free turning/walking; RLT/RLW; Real life turning/walking). As gait measures, we examined Lateral Step Deviation and a Compound measure of Spatial Variability, as these two gait measures were shown to have the highest sensitivity for ataxia severity ^5^. Average gait speed is provided as a general, i.e. non-ataxia specific measure of walking performance. Given are the correlations for healthy subjects (HC) as well as for all subjects with degenerative cerebellar diseases (DCD, consisting of preataxic and ataxic subjects). (*≡ p<0.05, **≡ p<0.01 Bonferroni-corrected, ***≡ p<0.001). Effect sizes of correlations are given using Spearman’s ρ. LVC: Lateral Velocity Change.

**Table 10.** Cross-sectional analysis of gait measures. Between-group differences in gait measures for healthy controls (HC), preataxic subjects (PRE) and ataxic subjects (ATX) in the different study conditions: Lab-based Walking (LBW), Supervised Free Walking (SFW) and Real Life Walking (RLW). Stars indicate significant differences between groups (*≡ p<0.05, **≡ p<0.01 Bonferroni-corrected, ***≡ p<0.001). KW denotes the result of the Kruskal-Wallis test. δ denotes effect sizes determined by Cliff’s delta.

| Parameter | Cond. | Descriptive Statistics |  |  | Statistical Test Results |  |  |  |  |  |  |
| --- | --- | --- | --- | --- | --- | --- | --- | --- | --- | --- | --- |
|  |  | HC | PRE | ATX | KW | HC vs PRE |  | HC vs CP |  | PRE vs CP |  |
| | | $\mu \pm \sigma$ | $\mu \pm \sigma$ | $\mu \pm \sigma$ | p | p | $\delta$ | p | $\delta$ | p | $\delta$ |
| Lateral Step Deviation | LBW | 2.61 ± 0.63 | 2.51 ± 0.37 | 3.72 ± 0.89 | <0.001*** | 1.00 | 0.01 | <0.001*** | 0.68 | 0.008** | 0.77 |
| | SFW | 3.86 ± 0.4 | 3.84 ± 0.33 | 4.43 ± 0.47 | <0.001*** | 0.833 | 0.06 | <0.001*** | 0.66 | 0.004** | 0.71 |
| | RLW | 3.81 ± 0.31 | 3.88 ± 0.27 | 4.31 ± 0.75 | 0.008** | 0.665 | 0.12 | 0.003** | 0.53 | 0.056 | 0.5 |
| Compound spatial variability | LBW | 0.18 ± 0.1 | 0.16 ± 0.09 | 0.37 ± 0.18 | <0.001*** | 0.809 | 0.07 | <0.001*** | 0.71 | 0.005** | 0.76 |
| | SFW | 0.24 ± 0.08 | 0.23 ± 0.05 | 0.34 ± 0.12 | 0.017** | 0.887 | 0.04 | 0.013* | 0.54 | 0.021** | 0.59 |
| | RLW | 0.21 ± 0.09 | 0.23 ± 0.07 | 0.34 ± 0.13 | 0.001*** | 0.51 | 0.17 | <0.001*** | 0.65 | 0.026** | 0.55 |
| Gait Speed | LBW | 1.33 ± 0.15 | 1.33 ± 0.14 | 1.32 ± 0.27 | 0.986 | 0.896 | 0.05 | 0.981 | 0.01 | 0.905 | 0.04 |
| | SFW | 1.34 ± 0.1 | 1.32 ± 0.09 | 1.21 ± 0.2 | 0.019* | 0.76 | 0.08 | 0.008** | 0.49 | 0.067 | 0.46 |
| | RLW | 1.28 ± 0.17 | 1.24 ± 0.14 | 1.11 ± 0.19 | 0.009** | 0.524 | 0.17 | 0.003** | 0.55 | 0.143 | 0.39 |

### Activities-Specific Balance Confidence (ABC) Scale and specific questions to confidence in turning movements

The Activities-specific Balance Confidence (ABC) Scale ^28^ is a 16-item questionnaire/survey. Each item is rated from 0% (no confidence) to 100% (complete confidence).

“How confident are you that you will not lose your balance or become unsteady when you …

1. walk around the house? %
2. walk up or down stairs? %
3. bend over and pick up a slipper from front of a closet floor? %
4. reach for a small can off a shelf at eye level? %
5. stand on tip toes and reach for something above your head? %
6. stand on a chair and reach for something? %
7. sweep the floor? %
8. walk outside the house to a car parked in the driveway?
9. get into or out of a car? %
10. walk across a parking lot to the mall? %
11. walk up or down a ramp? %
12. walk in a crowded mall where people rapidly walk past you? %
13. are bumped into by people as you walk through the mall? %
14. 14… step onto or off of an escalator while you are holding onto a railing? %
15. step onto or off an escalator while holding onto parcels such that you cannot hold onto the railing? %
16. walk outside on icy sidewalks? %

In addition to the validated questions of the ABC score, we asked the patients specific questions about their balance confidence in everyday life:

“ How confident are you that you will not lose your balance or become unsteady when you …

1. turn right/left from the corridor into a room (question - 90° Turns)
2. someone calls you from behind and you turn around (question - 180° Turns)

